# IMMREP25: Unseen Peptides

**DOI:** 10.64898/2026.03.30.715276

**Authors:** Eve Richardson, Yannick Jurriaan Maria Aarts, John A. Altin, Coos A. B. Baakman, Philip Bradley, Binbin Chen, Joakim Clifford, Manjima Dhar, Danielle Diepenbroek, Ethan Fast, Ragul Gowthaman, Jieling He, Vadim Karnaukhov, Dario F. Marzella, Pieter Meysman, Morten Nielsen, Jonas Birkelund Nilsson, Sebastian Nymann Deleuran, Farzaneh M. Parizi, Aurelien Pelissier, Brian G. Pierce, Maria Rodriguez Martinez, Dona Roran A R, Shayana Saravanakumar, Yanjun Shao, Nils Smit, Max Van Houcke, Gian Marco Visani, Yat-Tsai Richie Wan, Xiaowo Wang, Lawson Woods, Sander Wuyts, Chengkai Xiao, Li C. Xue, Justin Barton, Matthew Noakes, Damon H. May, Bjoern Peters, IMMREP25 Participant Consortium

## Abstract

T cell receptors (TCRs) can bind to peptides presented by MHC molecules (pMHC) as a first step to trigger a T cell response. Reliable approaches to predict TCR:pMHC binding would have broad applications in clinical diagnostics, therapeutics, and the fundamental understanding of molecular interactions. IMMREP is a community organized series of prediction contests that asks participants to predict TCR:pMHC binding on unpublished datasets. Previous iterations in 2022 and 2023 showed multiple approaches can predict TCR-pMHC binding with significant accuracy (median AUC_0.1≥0.7) for peptides where experimental data is available (“seen” peptides). In contrast, models did not outperform random guessing for peptides that have no such data available (“unseen” peptides). Here we report on the results of IMMREP25, which focused solely on unseen peptides in order to evaluate the cutting edge of the field. We received 126 named submissions predicting the specificity of 1,000 TCRs against twenty unseen peptides restricted by one of two MHC molecules (HLA-A*02:01 and HLA-B*40:01). The best performing methods showed a macro-AUC_0.1 of 0.60, significantly better than random, demonstrating significant advances in the field. The top performing methods incorporated structural modeling into their approach, indicating that especially for ‘unseen’ peptides, a structural understanding aids in the prediction of TCR:pMHC interactions. The results from this benchmark highlight the significant challenges remaining for TCR:pMHC predictions and will inform future method development.

## 1 Introduction

T cell receptors (TCRs) are expressed by T cells to scan ligands presented to them by major histocompatibility complex (MHC) molecules on the surfaces of other cells. Most of the MHC-presented ligands are peptides, and the peptide-MHC complex is referred to as pMHC. Each T cell has a unique TCR, generated through a complex stochastic process. T cells with TCRs that bind self-pMHC with high affinity are selected against during T cell maturation. Thus, binding of a TCR from a mature T cell to a pMHC signals the presence of a non-self-peptide and triggers a T cell response. Such T cell responses are crucial for the elimination of pathogens and cancer cells and understanding which TCRs can bind which pMHC provides a pathway for the generation of new diagnostic and therapeutic approaches in numerous diseases.

While methods for identifying TCR-pMHC pairs are increasing in throughput, these are still not accessible on the deep repertoire scale^1,2^. Reliable predictive models of TCR-pMHC binding hold enormous potential for providing new insights into targets recognized by TCRs sequenced via higher-throughput and cheaper methods^3^. The earliest predictive approaches rely on making mappings from large databases of TCRs of known pMHC specificity (such as VDJdb, mcPAS-TCR and the IEDB) to query TCRs using a look-up method, such as a kernel similarity over the hypervariable CDR3b^4–9^. More recent advancements in look-up methods include physicochemical and structural similarity calculated across all CDRs^10,11^. There have been successful attempts to improve on look-up methods through training machine learning models on a larger number of TCR features^12–24^. A limited number of approaches have used TCR-pMHC structure prediction directly^14,25–28^.

IMMREP is a series of prediction contests initiated in 2022 where contestants are invited to submit predictions of TCR-pMHC binding on novel test datasets^29,30^. In the first iteration (IMMREP22), contestants had to use the training data provided; in IMMREP23, training data was not proscribed, and multiple labs collaborated in producing a novel test dataset of 20 pMHC, including three pMHCs for which there were no TCRs in the public domain (unseen peptides). IMMREP23 revealed that for pMHCs for which there is a considerable body of positive TCRs, machine learning models such as ImmuneWatch DETECT, as well as deep learning frameworks such as NetTCR and MixTCRpred, had reasonable success^12,13,31^. However, these methods failed on unseen peptides, with no discernible generalizability advantage afforded by machine learning methods; this finding was echoed by a recent independent assessment of 21 prediction methods^32^.

As the next frontier in TCR-pMHC binding prediction, IMMREP25 tested contestants’ ability to make predictions on entirely unseen pMHCs. A novel dataset of 1,000 TCRs to 20 virally-derived peptides across two MHC molecules (HLA-A*02:01 and HLA-B*40:01) was expressly created for this purpose. Here, we describe the current state-of-play for prediction on unseen pMHCs, including a convergence on structural methods by the winning teams.

## 2 Methods

### 2.1 Evaluation dataset experimental methods

A novel dataset of 1,000 unique TCRs positive to one of 20 unseen pMHCs, with 50 TCRs per pMHC, was provided for the express purpose of the competition by Adaptive Biotechnologies. The evaluation dataset comprises a total of 10,000 records divided between 1,000 positive labels and 9,000 negative labels. These peptides were drawn from 2 MHC context, with 10 peptides coming from each of HLA-A*02:01 and HLA-B*40:01. The peptides themselves are all 9-mer epitopes from a variety of viral sources (**Table 1**) and all novel relative to those with previously published and collated T cell epitope datasets. Specifically, this dataset includes only peptides with little similarity to previously published epitopes as found in IEDB, VDJdb, McPAS or previously published MIRA data^7–9,33^. All peptides are 4 or more Levenshtein distance from any published peptide and share no substring longer than 5 consecutive amino acids with any published peptide.

**Table 1:**
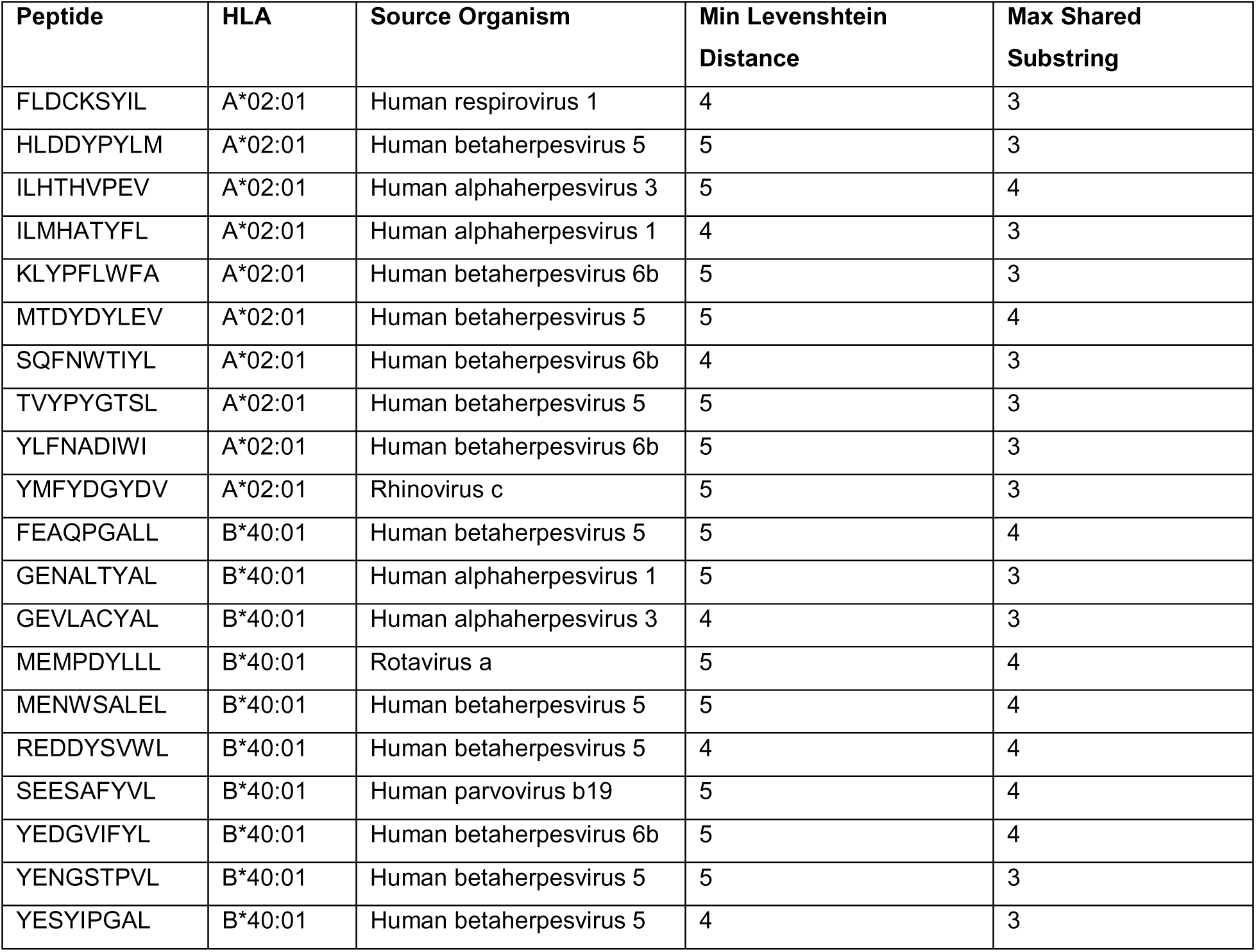
The 20 peptides represented in this dataset, described by their HLA context, source organism, minimum Levenshtein distance to any published epitope, and maximum shared substring with any published epitope.

PBMCs from HLA-genotyped donor leukopaks were isolated via negative selection for naïve, CD8+ T cells, and incubated with a pool of peptides for T cell expansion. A portion of this expanded T cell population was restimulated with the entire pool of peptides and input into a pairSEQ experiment to determine TCRA-TCRB pairings as described in Howie et al ^34^. The remainder of the peptide pool expanded T cell sample was input into a MIRA (Multiplex Identification of T cell Receptor-Antigen specificity) experiment from which we determined TCRB-peptide specificities as described in Klinger et al (2015), Nolan et al (2024) (experimental methods) and Snyder et al (2025) (statistical methods)^1,33,35^.

From the TCRA-TCRB pairings (from pairSEQ) and TCRB-peptide specificities (MIRA), we constructed complete TCR-peptide information. Finally, we defined the MHC context using peptide-MHC binding prediction from NetMHCpan4.1 and limiting it to data in which the purported presenting MHC for the peptide was also present in the source donor’s HLA genotype^36^. In combination, this process generated the positive-label TCR-pMHC data.

For the negative-label TCR-pMHC data, we took advantage of the multiplexed nature of the MIRA assay **Figure 1**). In MIRA, TCRB clonotypes are assessed for activation against all peptide pools in the experiment and only reported as positive for a given peptide when their set of activated pools corresponds to the unique occupancy pattern of that peptide. Consequently, any TCRB reported as positive must not activate in response to any other peptide in the experiment, as this would result in activation in additional pools and disqualify it from being labeled as a positive. Since all 20 peptides in the evaluation dataset were assessed simultaneously in each MIRA experiment, every positive TCR is a valid negative for the remaining peptides.

### 2.2 Evaluation dataset processing

For each MHC, the dataset includes 50 TCRs which were experimentally determined to be specific for that pMHC by MIRA and pairSEQ^1,34^ (Section 2.1). Each of these pMHC-specific TCRs was determined to not be specific for any of the other 19 pMHCs in the dataset. The complete labeled dataset consists of these 1,000 positive TCR-pMHC combinations and the 9,000 negative TCR-pMHC combinations formed by matching each TCR to all alternative peptides within its ground truth MHC context. Negative records consisting of TCRs against peptides outside their ground truth MHC context were not included in the dataset. These combinations were then randomly downsampled to 8,938 records so that contestants could not assume an even distribution of TCRs per pMHC.

The records for two HLA-A*02:01-presented peptides were used for the public leaderboard, corresponding to 175 entries. Records from the remaining 18 pMHCs form the private dataset used for evaluation, corresponding to 7,344 records. The remaining 1,419 records correspond to TCRs positive to a public peptide and negative to a private peptide, or vice versa. These were excluded from both public and private leaderboard calculations to prevent any validation to holdout leakage.

### 2.3 Prediction method summaries

#### Bradley methods (contributed by Philip Bradley)

For the AF3TD predictions, we used a version of the TCRdock pipeline, updated to use Alphafold3 (AF3) and MSAs, to dock each TCR onto all potential partner peptides and ranked the TCR:pMHC pairings based on Alphafold metrics^25,37^. TCRdock is an Alphafold-based pipeline that uses inter-chain template geometries to encourage canonical TCR:pMHC docking^25^. The matrix (500 TCR rows by 10 peptide columns) of raw AF3 metric values (either ipTM or pLDDT (top submission) averaged over peptide and CDR loops) was normalized by subtracting the row and column means.

For the cluster-mode submissions, matrix rows were clustered by TCR sequence, and row values were averaged over the clusters, upweighted by the square root of the cluster size (to counteract the tendency of averaging to smear things out). In the “small cluster” submission, clusters were formed by single linkage with a TCRdist threshold of 120^5^. In the “big cluster” submission, the TCRs were divided into 10 equal groups of 50 by hierarchical TCRdist clustering.

#### Altin method (TCR-TriFold) (contributed by Lawson Woods and John Altin)

AF3 was run with MSAs and templates turned on^37^. All triads (TCR-pMHC combinations) were run with five seeds (1–5), a single diffusion sample, and 10 recycles. For each triad, we used AF3 to predict the structure (from sequence) of the protein complex formed by its 5 constituent protein chains: peptide, MHCα, MHCβ, TCRα, TCRβ. Based on prior training of features on data from the IEDB, we scored the outputs using the mean Predicted Alignment Error (PAE) of the peptide:TCR interface^7^.

#### Pierce method (contributed by Ragul Gowthaman, Shayana Saravanakumar and Brian G. Pierce)

AF3 was used to model TCR-pMHC complexes, utilizing the reduced MSA database from TCRmodel2 for computational efficiency^37,38^. Interaction scores were obtained by averaging TCR-pMHC inter-chain predicted alignment error (PAE) scores for the top-ranked model, then linearly scaled (0-1) with a PAE ceiling of 10 Å. Any modeled interaction with fewer than five peptide residues contacted by the TCR was assigned a score of 0. For a separate submission, we additionally clustered the TCR sequences at 96% identity and assigned high confidence pMHC targets (score >0.85) based on co-clustered TCRs.

#### Gowthaman method (contributed by Ragul Gowthaman, Shayana Saravanakumar and Brian G. Pierce)

We employed XGBoost classifiers to distinguish TCR binders from non-binders using curated TCR-peptide-MHC features. Positive training data comprised 384 high-confidence TCR-pMHC sequences from VDJdb complexes with VDJdb confidence score 3, while the negative set included 404 randomly paired TCR and pMHC sequences from PDB structures and VDJdb complexes^9^. Two XGBoost models were evaluated: one using only AF3 metrics (ipTM, pTM, ranking score, multimer confidence score, and average peptide pLDDT) and another combining these metrics with 1280-dimensional ESM-2 embeddings of CDR3α, CDR3β, and the peptide^39^. The AF3 metrics-only model was the top-performing submission for this method.

#### Wang methods (contributed by Jieling He and Xiaowo Wang)

We used AF3 to predict the complex structure and directly took the model-reported iPTM (top submission) or PAE score (computed as exp(-mean_pae) and then normalized) as the binding prediction score^37^. We did not apply additional learning, calibration, or post-processing beyond the AF3 run settings. Since generating MSAs is time-consuming, we made a small optimization: we did not compute MSAs for peptides; for MHC sequences (limited to a few specific variants) we precomputed MSAs; and for TCR sequences we first computed MSAs for a small subset, collected those results into a new database, and then generated MSAs for subsequent sequences by searching against this new database.

#### TAPIR3 (contributed by Binbin Chen, Manjima Dhar and Ethan Fast)

TAPIR3 is a sequence-based model based on fine-tuning ESM (PMID: 36927031) to predict structure-derived metrics (pTM, ipTM, iPAE) generated by Chai-1^39,40^. Training incorporated known TCR-pMHC binding pairs from IEDB, the Kaggle training set, and internal datasets (Vcreate), with additional loss weighting applied to these positive interactions. Input representations for paired TCR and pMHC sequences followed encoding schemes similar to those described in TAPIR^19^. This knowledge distillation approach enabled rapid inference while leveraging structural information from the protein folding model.

#### Nielsen (contributed by Joakim Clifford, Sebastian Nymann Deleuran, Yat-Tsai Richie Wan, Jonas Birkelund Nilsson and Morten Nielsen)

For each TCR–pMHC, Chai-1 (pip package version 0.6.0) is run^40^. 20 structures are predicted for each TCR–pMHC (using four seeds, each with five diffusion samples). Each predicted structure is assigned a score (the default Chai-1 aggregate score; 0.2 pTM + 0.8 ipTM). For each TCR-pMHC, the final score is obtained by max pooling across the 20 structures.

For the cluster method, TCR sequences are clustered using TCRCluster with the “Two Stages (No Triplet)” setting^41^. For each TCR cluster: 1. Apply z-score normalization. 2. Identify the max-scoring TCR–pMHC in the cluster. 3. Identify the peptide of the max-scoring TCR–pMHC. 4. Identify other confident peptides belonging to TCR–pMHCs scoring within 2.5% of the max-scoring TCR–pMHC. 5. Define confident peptides as: [max-scoring peptide] + [other confident peptides] 6. For each confident peptide, transfer the max score from Step 2 to other TCR–pMHCs with the same peptide. Chai1-TCR-clus: Uniformly downweight (*0.25) scores for TCR–pMHCs with non-confident peptides. Chai1-TCR-clust2 Downweight (*0.25 × best_raw_score) scores for TCR–pMHCs with non-confident peptides. ‘best_raw_score’ is the score from Step 2 before z-score normalization.

#### Xiao (contributed by Chengkai Xiao)

We predicted the structure of the TCR-pMHC complex using the Chai-1 model with ESM embeddings as input^39,40^. The final prediction was determined by calculating the average Chai-1 aggregate score across five models.

#### NetTCR-struc (Joakim Clifford, Sebastian Nymann Deleuran, Yat-Tsai Richie Wan, Jonas Birkelund Nilsson and Morten Nielsen)

AlphaFold 2.3 was run as described in Deleuran et al with four seeds, generating 20 structures per TCR-pMHC (each seed producing five predictions)^14,42^. Each predicted structure was assigned a score by NetTCR-Struc, yielding 20 scores per TCR-pMHC. For each TCR-pMHC, the final score was obtained by maxpooling. For NetTCR-Struc cluster methods, the same clustering algorithm as described in the Nielsen Chai-1 method was used.

#### Immunewatch (contributed by Max Van Houcke, Pieter Meysman and Sander Wuyts)

For each TCR-pMHC combination, Boltz-1 was used to infer a 3D structure confidence score and ImmuneWatch DETECT was run on motif mode to infer a sequence-based confidence score^43^. Both scores were normalised using a per-peptide z-normalisation and the average score of both tools was used as a submission. Input alignments for Boltz 1 were generated using a local installation of localcolabfold^44^. Boltz 1 with the pre-aligned sequences was run on the VSC (Flemish Supercomputer Center), funded by the Research Foundation Flanders (FWO) and the Flemish Government.

#### Visani (contributed by Gian Marco Visani)

TCR-pMHC structures were predicted with TCRdock (a version of AF2 optimized to perform better at folding TCR-pMHC complexes)^25^. PAE computed between TCR and pMHC chains was used as a raw score, which was then normalized across peptides and TCRs (independently for each MHC allele) by subtracting the mean score. This method corresponds to the top submission for this team.

An additional method, with significantly indistinguishable performance from the described TCRdock method, used HERMES to compute interaction scores^28^. HERMES was pretrained to predict amino acid identity of a masked residue given its atomic environment, and then fine-tuned such that its output log-probability difference between two amino acids matches experimental ΔΔG stability measurements. Following Visani et al (2025), we computed raw TCR-pMHC interaction scores by computing the log-probability of the observed peptide sequence in the folded structure^45^.

#### TCRen (contributed by Vadim Karnaukhov)

Predictions were based on TCR–peptide interaction energies computed using the TCRen statistical potential derived from amino acid contact preferences in TCR–pMHC crystal structures^27^. For each test pair, a TCR–peptide–MHC complex was modeled using AlphaFold3-TCRDock, and TCR–peptide interaction score was calculated as the sum of TCRen values for all TCR–peptide pairwise residue contacts (≤5Å) in the predicted structure^25,37^. Scores were normalized by subtracting per-epitope and per-TCR means. TCR sequences were clustered using TCRdist (20 clusters of 50 TCRs), and final predictions combined raw and cluster-averaged scores^5^. Notably, AlphaFold3 confidence metrics were not used for predictions.

#### STAPLER (contributed by Aurélien Pélissier, Yanjun Shao and María Rodríguez Martínez)

Binding predictions were run for every possible peptide-TCR pair with STAPLER according to instructions on GitHub (https://github.com/schumacherlab/STAPLER), then prediction outputs were grouped into 20 equal-sized clusters using constrained K-means to ensure a balanced distribution of TCRs per group^46^. These hard assignments were finally converted into continuous ranking scores by a k-Nearest Neighbors (k-NN) model, which calculates the binding probability based on the cluster membership of a TCR’s 15 closest neighbors.

#### SwiftTCR (contributed by Yannick Jurriaan Maria Aarts, Coos Baakman, Danielle Diepenbroek, Dario F. Marzella, Farzaneh M. Parizi, Dona Roran A R and Li C. Xue)

SwiftTCR requires TCR and pMHC structures as input^47^. We used TCRmodel2 to generate five TCR–pMHC models per case, then re-docked them with SwiftTCR and clustered the poses using a 9 Å i-rmsd cutoff^38^. TCRs were ranked by the largest cluster size, assuming larger clusters indicate more plausible binding. We also performed HADDOCK water refinement on TCRmodel2 to obtain HADDOCK scores^48,49^. The final ranking combined the TCRmodel2 confidence score, total and electrostatic energies, normalized cluster size, and HADDOCK score, each individually min–max scaled and summed.

#### HADDOCK (contributed by Yannick Jurriaan Maria Aarts, Coos Baakman, Danielle Diepenbroek, Dario F. Marzella, Farzaneh M. Parizi, Dona Roran A R and Li C. Xue)

We used TCRmodel2 to generate TCR–pMHC complex models. Because HADDOCK refinement is computationally expensive, we first filtered out obviously incorrect models (e.g., those in which the TCR is not positioned over the top surface of the pMHC). We then performed a HADDOCK water refinement step on the remaining models^48,49^. The resulting HADDOCK total energy, electrostatic energy, and TCRmodel2 confidence score were each normalized between 0 and 1, summed, normalized again and used to rank the final models.

### 2.4 Evaluation metrics and methods

We calculate a macro or average AUC as per previous competitions, specifically the early retrieval ROC-AUC with maximum FPR set to 10%^29,30,50^. The AUC_0.1 is calculated per pMHC and averaged across the private set of pMHCs (n=18) to produce a macro score. ROC-AUC and AUC_0.1 are calculated using scikit-learn’s roc_auc_score function, and PR-AUC calculated using average_precision_score^51^.

## 3 Results

IMMREP25 was designed as a test of TCR-pMHC prediction methods on unseen peptides. To achieve this, we created a dataset of TCRs with positive and negative labels for 20 unseen pMHC molecules totaling 8,938 records (**Methods Section 2.1**). We established the macro-AUC_0.1 as the success metric by which competitors were ranked.

The competition was hosted on Kaggle (https://www.kaggle.com/competitions/immrep25) and was open for 21 days between March 2nd and March 23rd 2025, with one submission permitted per day per user. There was no restriction on the training data that competitors could use. Records from two pMHC molecules were used to evaluate models and report their performance in the public leaderboard as the competition progressed (public pMHCs). The remainder were used to evaluate the models and produce a private leaderboard at the completion of the competition; as contributors made multiple submissions per team, contributors could select their preferred submission for private evaluation.

While 73.0% of submissions exceeded random performance of 0.50, 50.0% of submissions fell in the range of 0.50<x<=0.52, while just 10.3% of submissions exceeded 0.55 (**Figure 2A**). Predictive methods with AUC_0.1 values exceeding 0.52, are described in **Table 2** (taking the top method per team, where multiple were submitted). The per-peptide score AUC_0.1 distribution of these methods is shown in **Figure 2B** (with the micro-ROC curves and associated AUCs in **Supplementary Figure 1**).

**Table 2:**
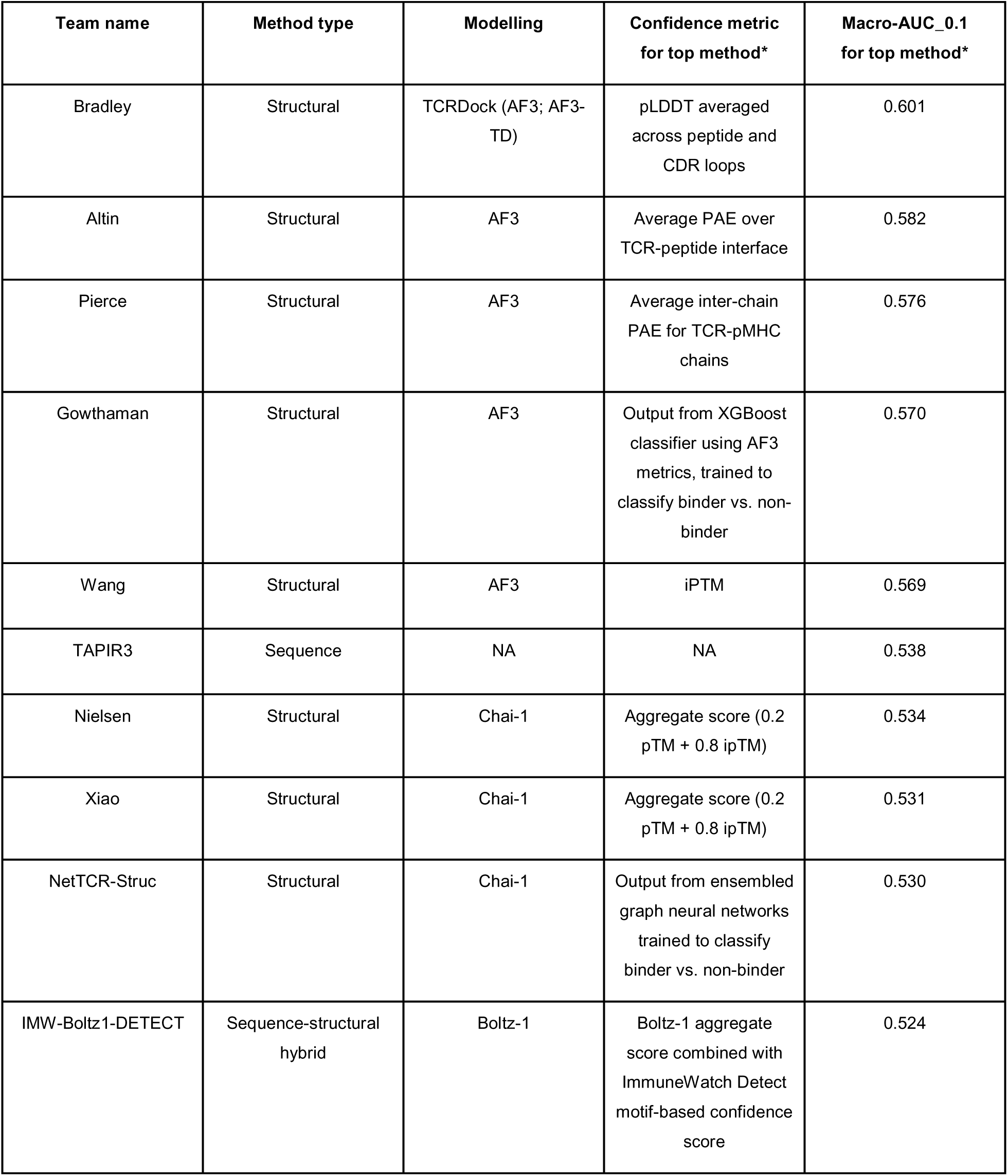
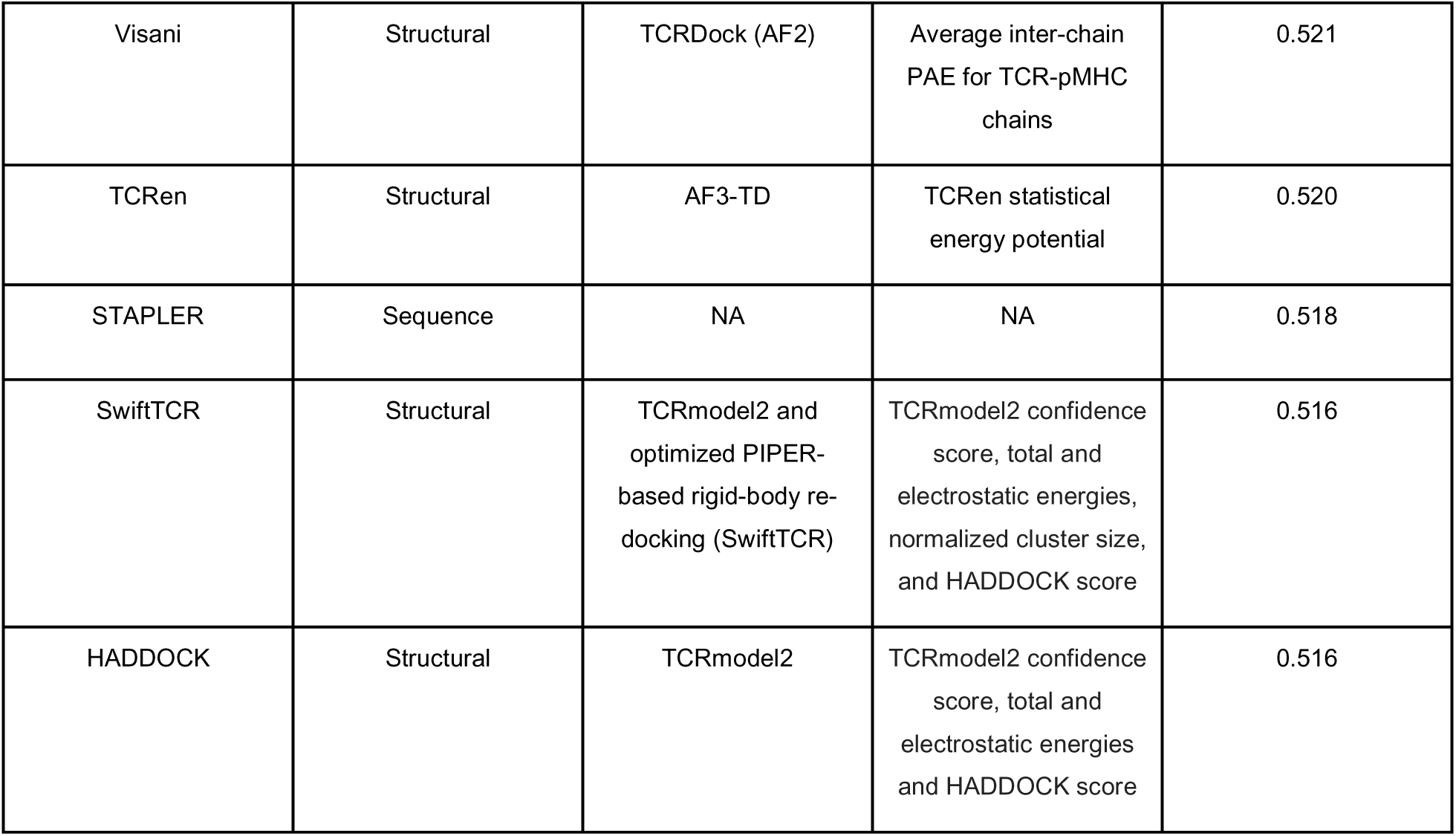
structural methods were well-represented among the methods with macro-AUC_0.1 ≥ 0.52 (only the top method per team is described). There was a further distinction between methods that used AF3 for TCR-pMHC modelling and alternatives such as Chai-1 and Boltz-1. *Methods details and scores are shown for the top submission for the team in question, as multiple teams submitted closely-related methods. **Supplementary Figure 3** and **4** show performance across all models submitted, and per-peptide and macro scores are found in **Supplementary File 1**.

Ranking by the macro-AUC varies slightly (ρ = 0.83) (**Supplementary Figure 2A**). The PR-AUC ranking likewise is largely concordant with the macro-AUC_0.1 (ρ = 0.92) (**Supplementary Figure 2B**). The full leaderboard score distribution for AUC_0.1 and AUC are shown in **Supplementary Figure 3** and **Supplementary Figure 4**.

There was significant heterogeneity between peptides and HLA molecules: predictions on the HLA-A*02:01-presented peptides were significantly more successful than HLA-B*40:01 on the whole and in the highlighted methods (p = 2.9e-9, t-test; **Supplementary Figure 5**). This likely reflects the difference in the HLA molecule distribution in training data (both TCR-specific training data and in the PDB): VDJdb and the IEDB contain 8,184 and 5,488 TCRs for HLA-A*02:01 respectively, and 46 and 50 TCRs for HLA-B*40:01. TCR3D lists 396 HLA-A*02 pMHC complexes (of which 265 are HLA-A*02:01; 129 are TCR-bound), and just a single HLA-B*40:01 pMHC complex (with none in complex with TCR) in the PDB^52,53^.

Throughout the course of the competition, some competitors shared that they were utilizing the assumed underlying structure of the dataset (**Supplementary Figure 6**) and that this substantially improved their performance in the public leaderboard. With the concern that this would complicate the interpretation of model skill, the organizers requested that subsequent to March 19th, competitors indicate whether their method utilized this structure via clustering. For seven methods which provided interpretable “raw” vs. “clustered” methods, cluster methods indeed provided a significant improvement on 2 pMHCs comprising the public leaderboard, but did not appear to provide a systematic advantage across the larger set of 18 pMHCs comprising the private leaderboard. Across such methods, there were no substantial improvements in macro-AUC_0.1 (**Figure 3**).

**Table 2** is almost entirely populated by structural methods. The top sequence-input model, TAPIR3, exceeded the performance of multiple structural methods: this model uses implicit structural information, as an ESM-based Chai-1 confidence metric distillation model, and impressively, outperformed the native Chai-1 ranking methods (Nielsen and Xiao). The only completely sequence-based method was STAPLER (macro-AUC_0.1 = 0.518). The remainder of published sequence-based methods which were assayed (EPACT, MINT, TULIP, TEIM, MixTCR-pred and NetTCR2.2) achieved AUC_0.1 values ranging from 0.500 to 0.510 confirming previous findings that such models exhibit poor performance on unseen peptides (**Supplementary Table 1**).

## 3 Discussion

IMMREP25 saw current TCR-pMHC binding prediction methods pitted against the challenging scenario of pMHC molecules for which there are no known TCRs in the public domain. The previous IMMREP23 competition showed that for three unseen pMHCs, no prediction methods succeeded without taking advantage of biases in the test dataset construction. The novelty of this iteration of the competition was its exclusive focus on unseen peptides, enabled by a novel dataset of 1,000 TCRs targeting 20 pMHC molecules.

In the intervening time since IMMREP23, there have been significant advances in protein complex structure prediction resulting in changes in approaches to TCR-pMHC binding prediction. TCR-pMHC binding prediction methods increasingly leverage protein language model and structural representations (such as inverse folding model representations) for featurization^14,54–56^. 2024 saw the release of details describing AlphaFold3, subsequent open-source models reproducing this method (Chai-1 and Boltz-1) and in late 2024, the release of AlphaFold3’s weights^37,40,43^.

The results from IMMREP25 both attest to the difficulty of the task and validate the shift in the TCR:pMHC binding prediction landscape towards next-generation structural methods. Most competitors failed to significantly outperform random: here we described the 15 methods with macro-AUC_0.1 values equal to or exceeding 0.52, of which two were sequence-based (TAPIR3 and STAPLER; STAPLER is arguably the only truly sequence-based model, as TAPIR3 is a sequence-input Chai-1 distillation model). A handful of methods significantly exceeded this selected threshold, with scores between 0.57 and 0.60. These highly successful methods all converged on a strategy of modelling TCR-pMHC complexes with AF3 and then using various native scoring methods (pLDDT (Bradley method), iPTM (Wang method) or PAE (Pierce and Altin methods)). Authors also explored TCR cluster-based pooling functions for a subset of these methods.

The top method, referred to as the Bradley method, used a modified AF3 version (AF3-TD) which encourages canonical TCR-pMHC geometries; in conjunction with AF3’s native confidence methods, this model outperformed the remainder of the AF3 methods, particularly in the macro-AUC. Another team used AF3-TD for TCR-pMHC complex prediction but combined it with a TCR-specific scoring function based on TCR-pMHC residue contact frequencies (TCRen), which significantly reduced performance across all metrics. While these methods differed additionally in clustering and normalization strategies, our overall survey of methods suggests that while TCR-pMHC specific knowledge can be meaningfully incorporated during complex prediction, there is not evidence that existing TCR-pMHC-specific scoring functions can outperform AF3’s confidence metrics.

IMMREP25 indicates that structural methods are the current leading method for identifying TCRs binding to unseen pMHCs. Several distinct structure-based methods demonstrate significantly non-random performance on this task, while also falling far short of perfect prediction. The top performing methods used a variety of natively-supplied confidence metrics from AF3. Beyond raw performance, a complication to the applicability of structure-based methods is that they are computationally resource-intensive, which limits their utility to limited sets of TCRs against a small number of possible pMHC molecules. This would exclude the bulk repertoire scale and where there are many TCRs and many possible pMHC molecules. There is therefore substantial scope for improvement of these methods, such as through distillation approaches like TAPIR3, which we envisage will be reflected in IMMREP26.

## Data availability

The evaluation dataset is provided for general use under an MIT License and can be downloaded from DOI: 10.6084/m9.figshare.30491582.

---

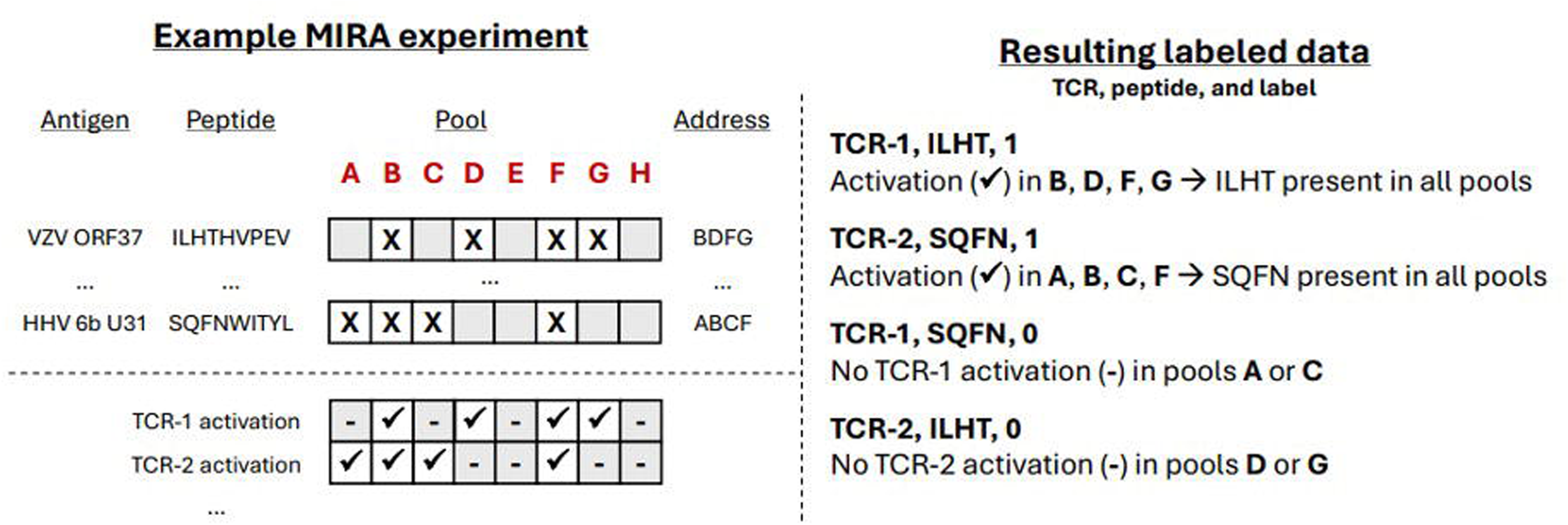

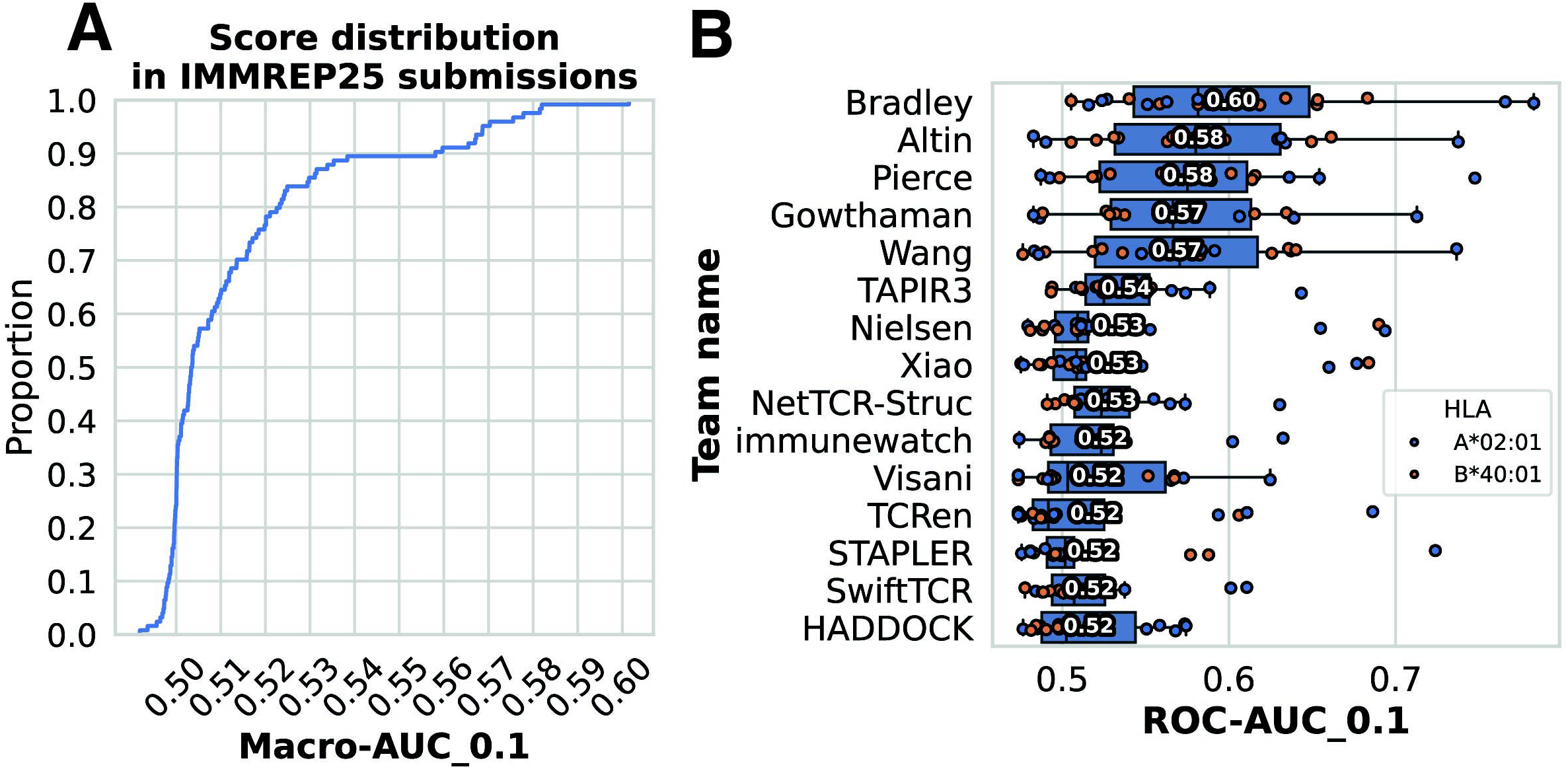

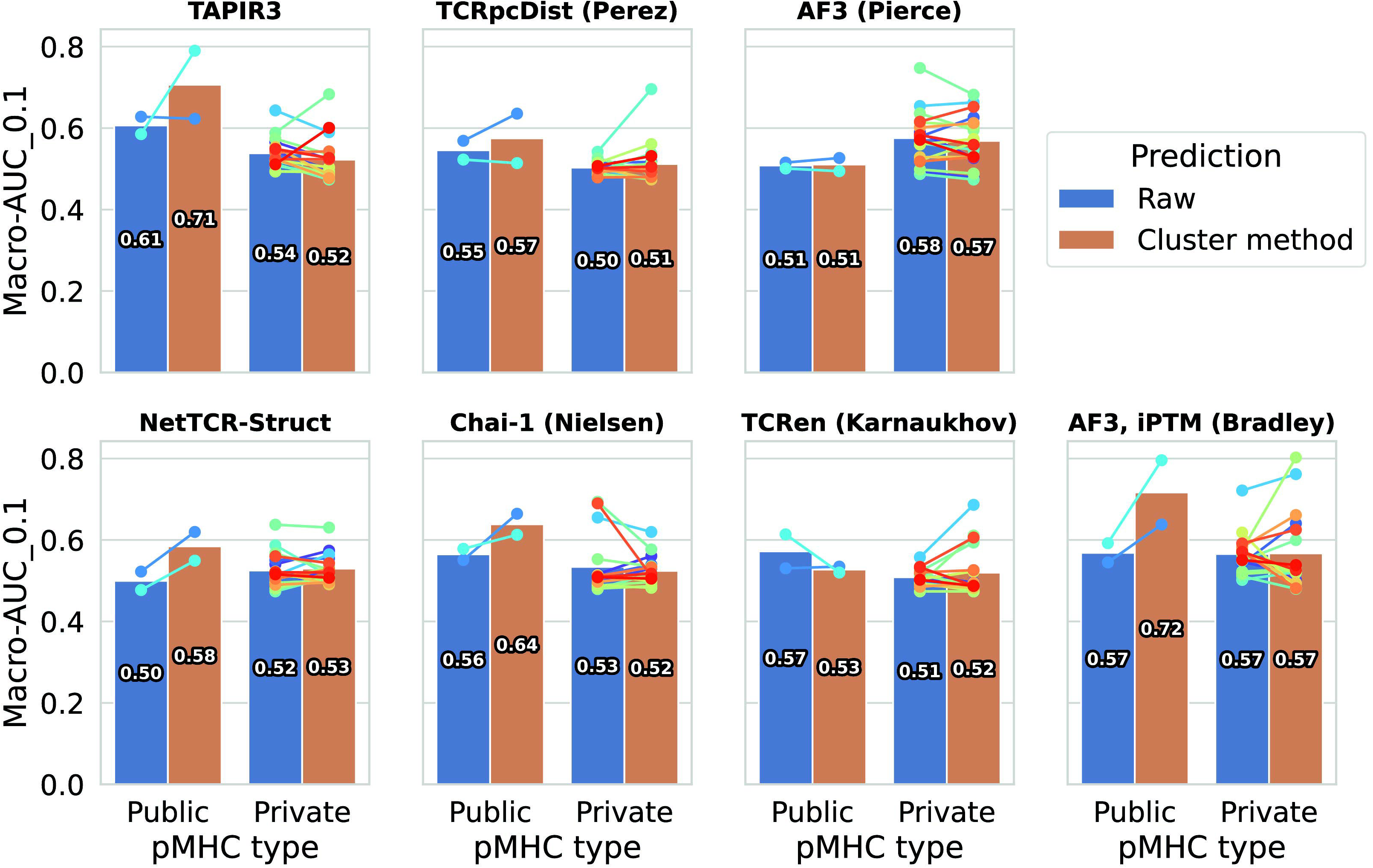

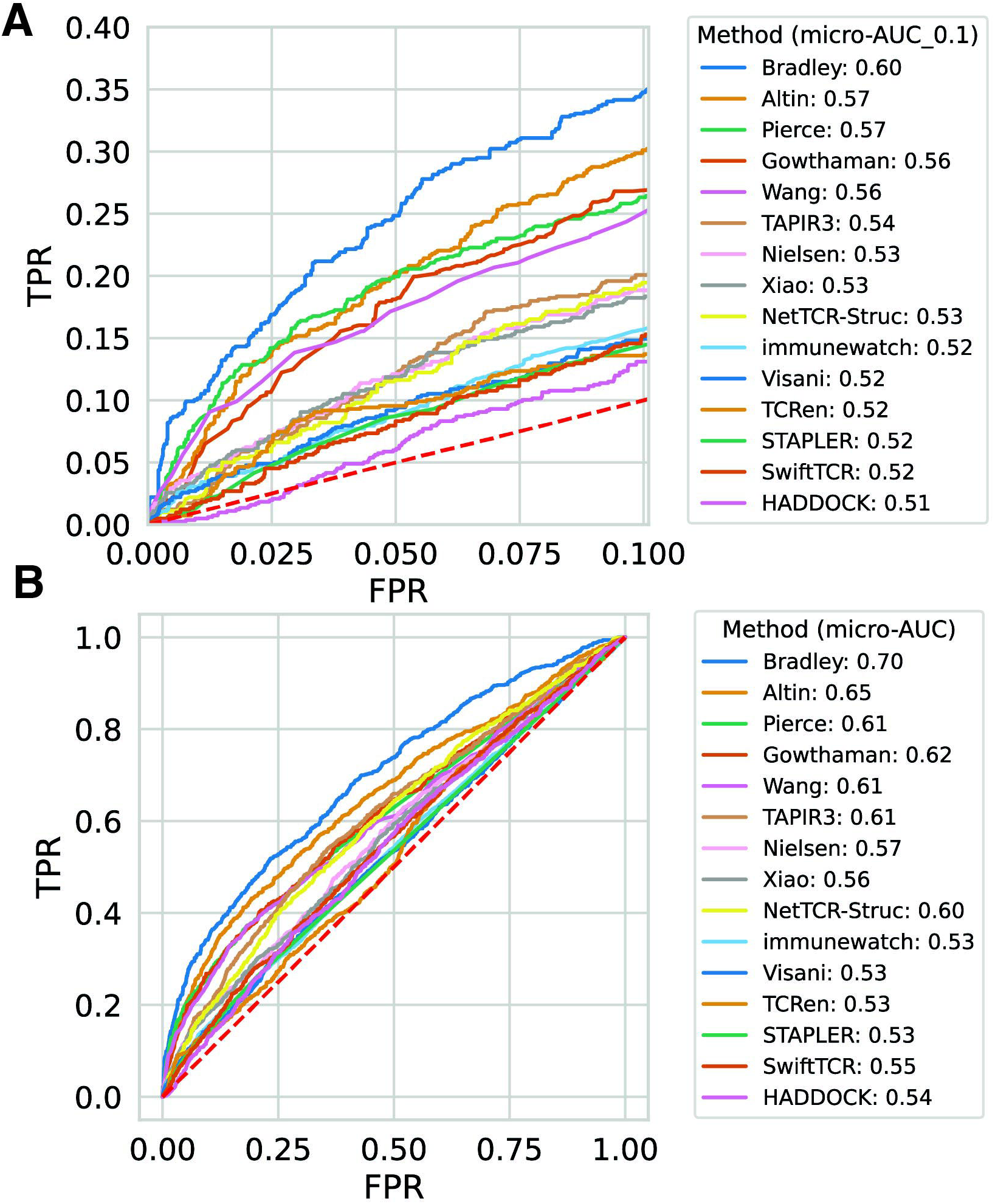

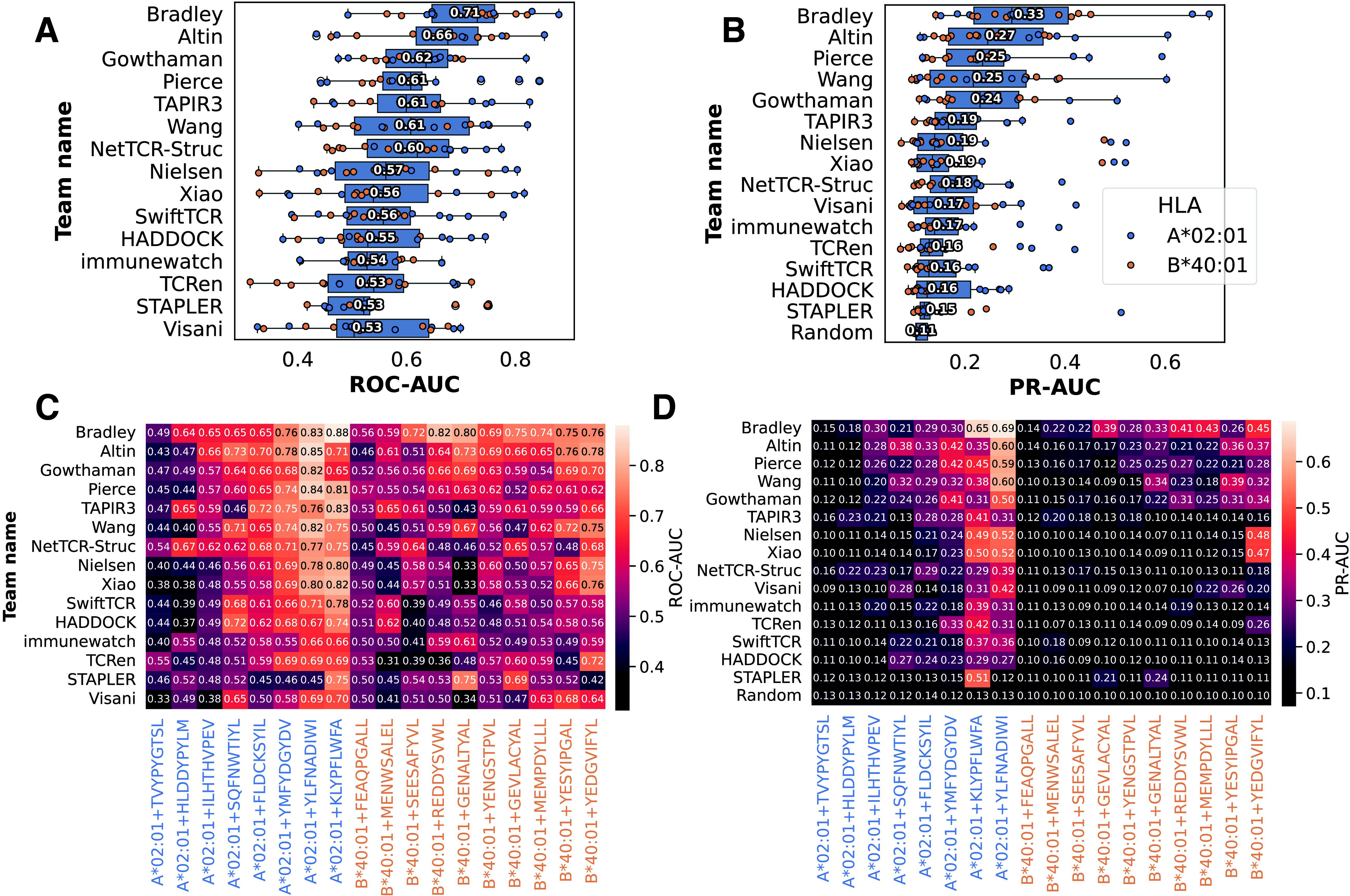

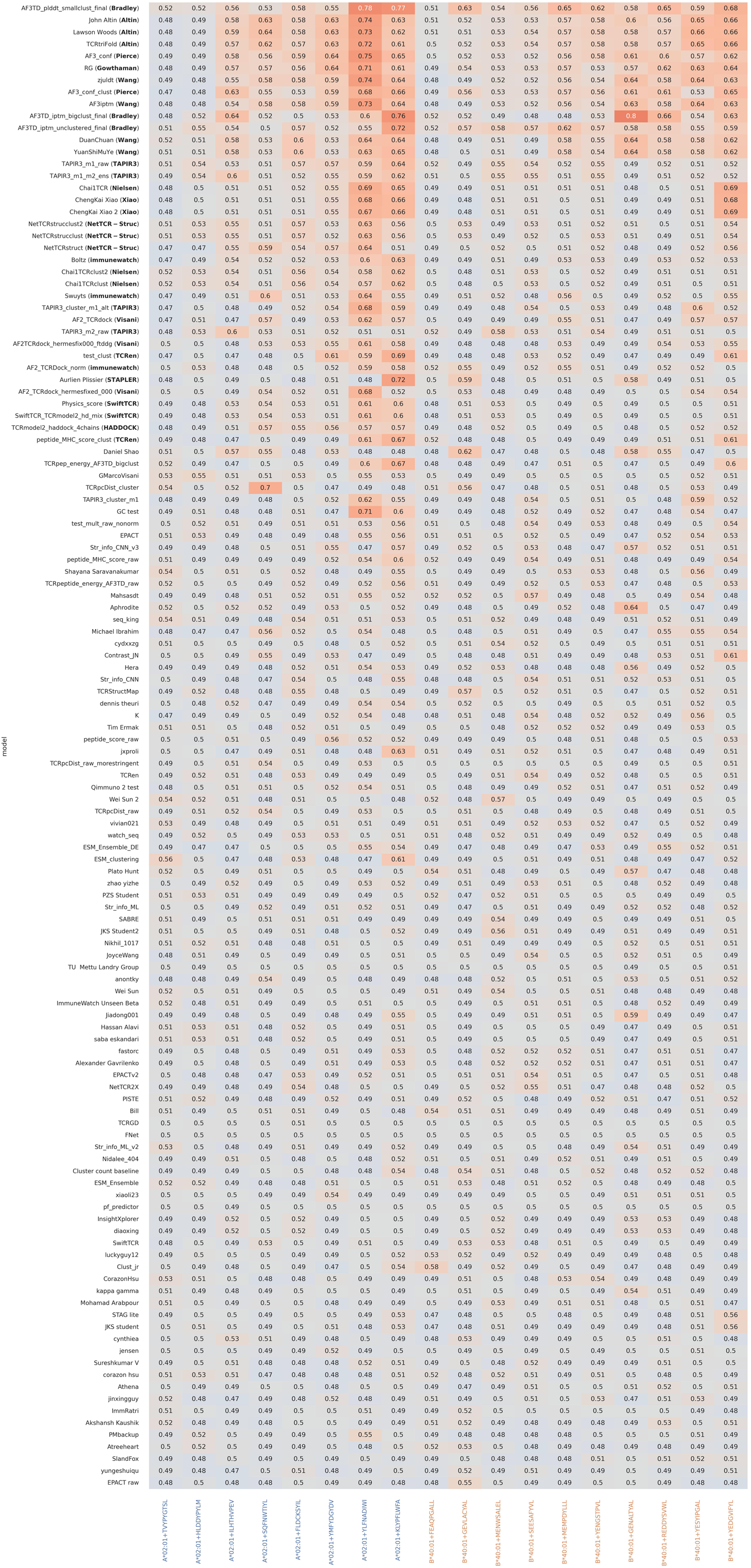

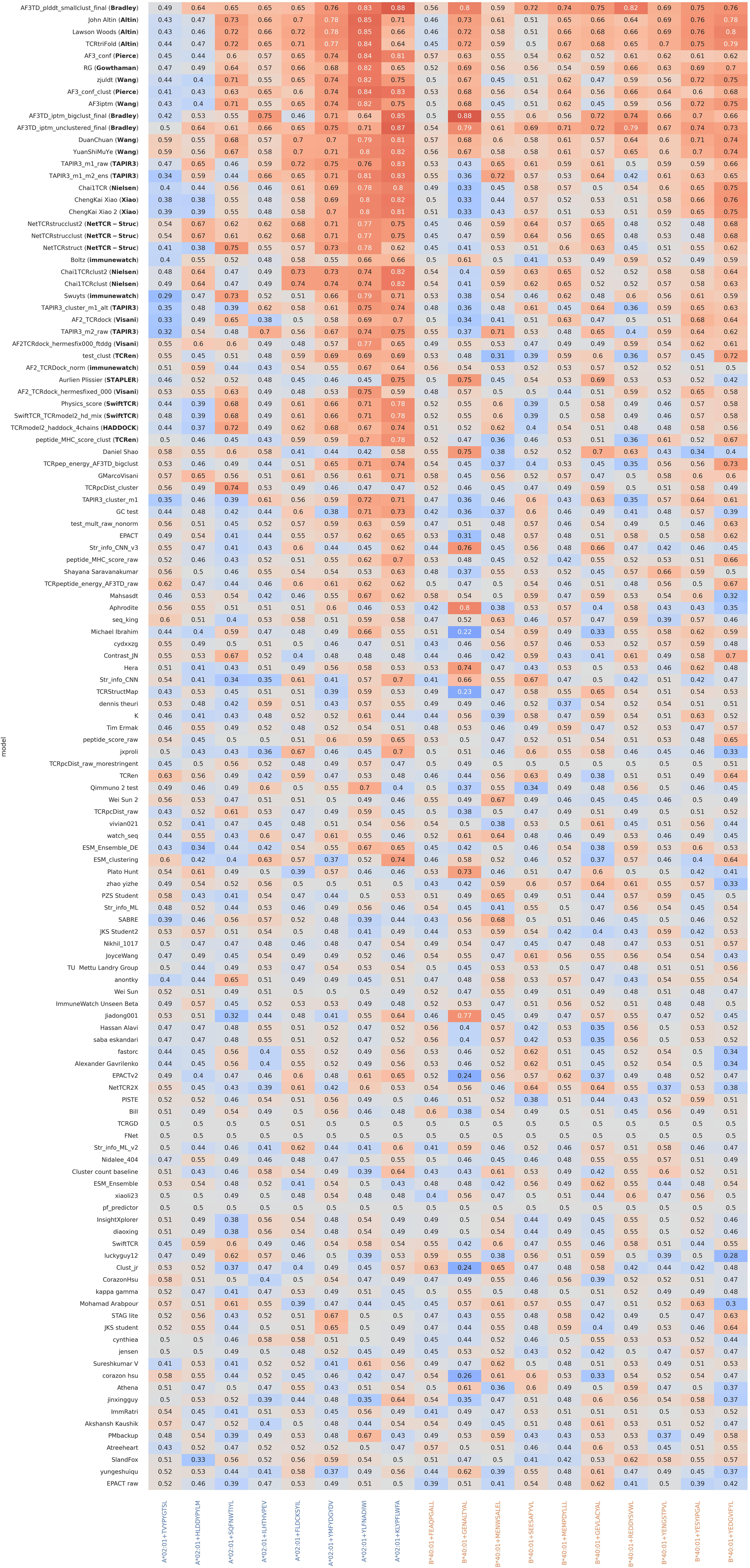

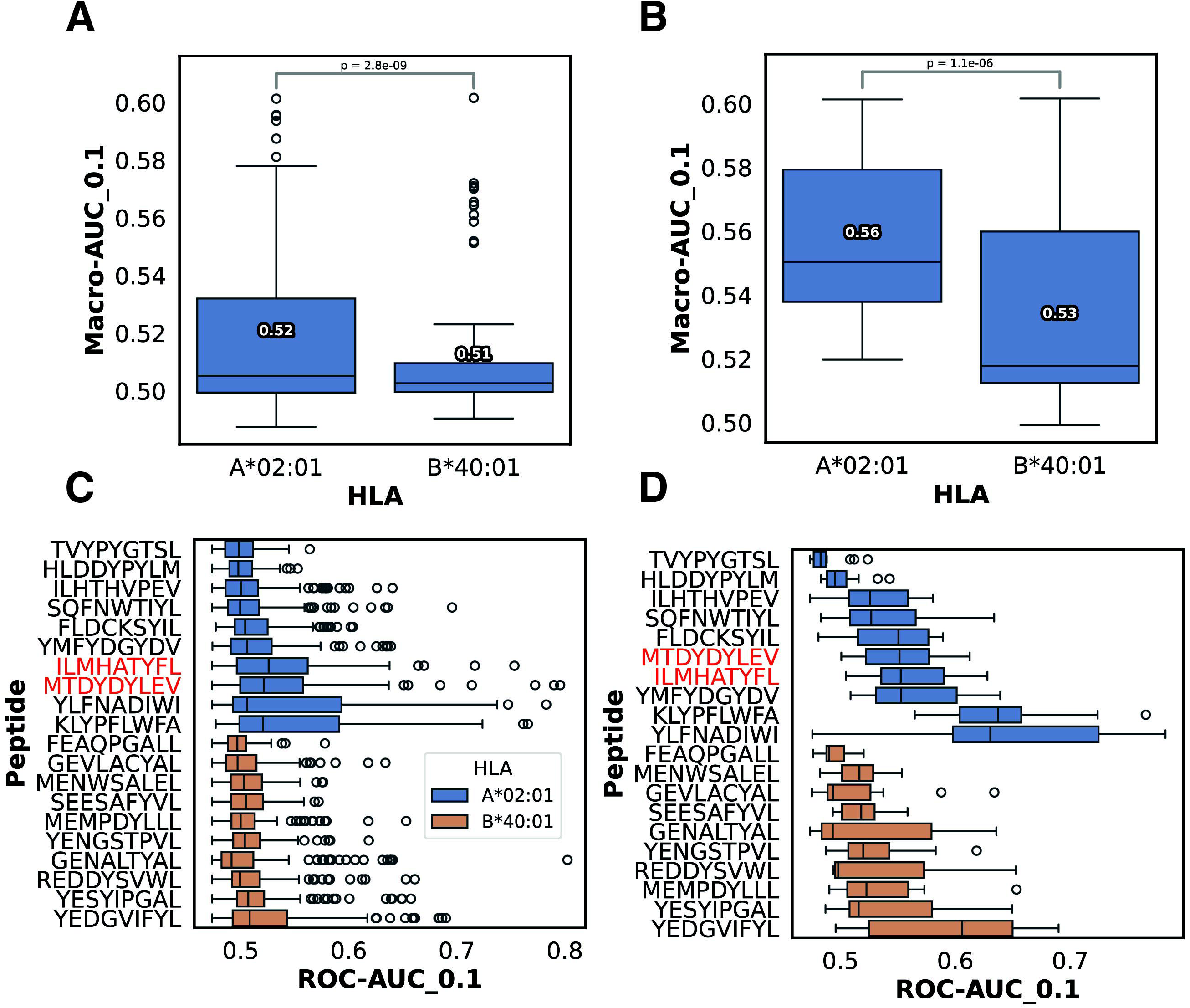

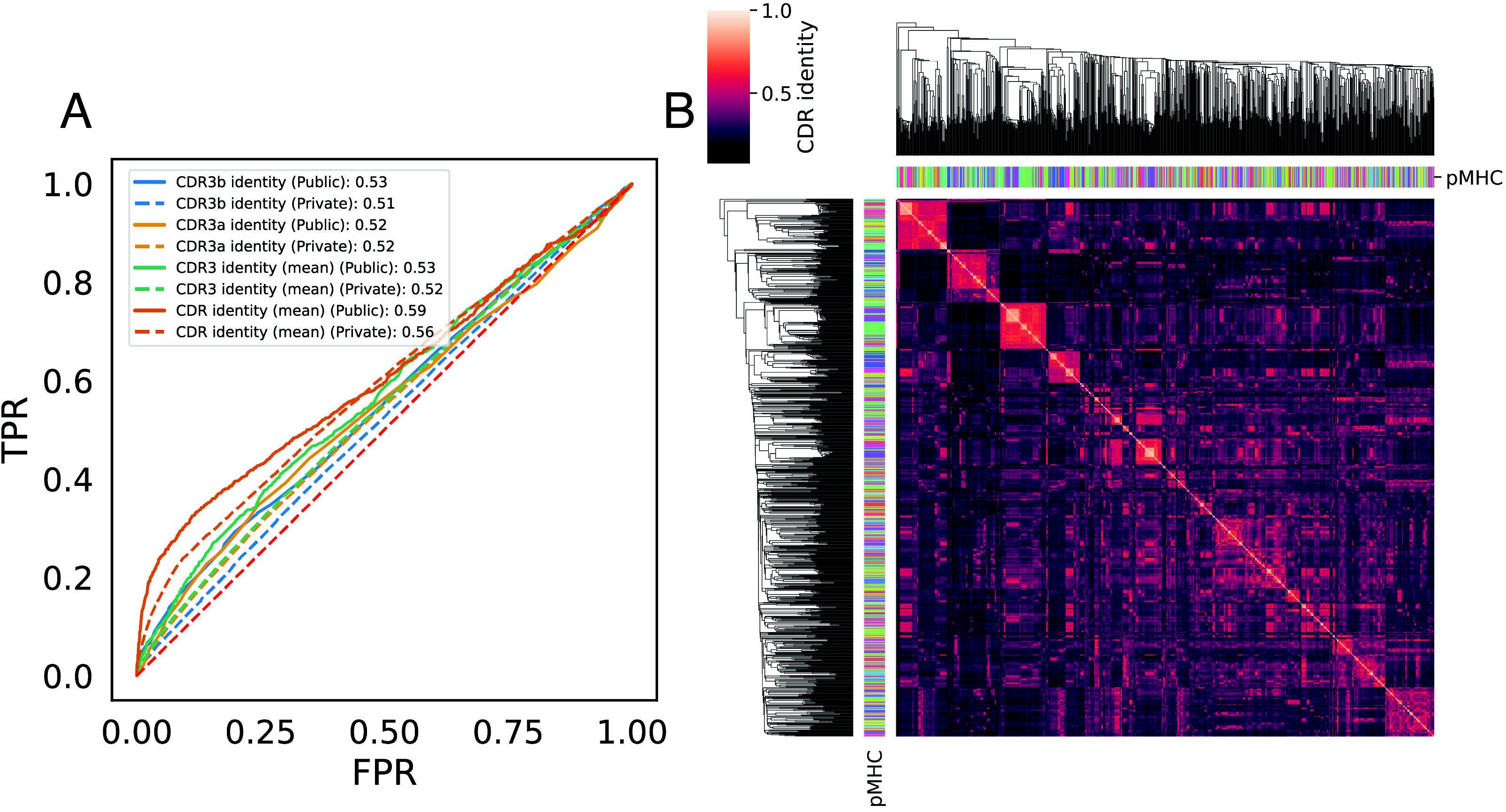

## Notes

https://figshare.com/articles/dataset/A_Public_Benchmark_for_TCR-pMHC_Prediction/30491582

### Competing Interest Statement

The authors have declared no competing interest.

