## Supplementary Information for "IMMREP25: Unseen Peptides"

### Supplementary Information
### IMMREP25: Unseen Peptides

Eve Richardson[1], Yannick Jurriaan Maria Aarts[2], John A. Altin[3], Coos A. B. Baakman[4], Philip Bradley[5], Binbin Chen[6], Joakim Clifford[7], Manjima Dhar[6], Danielle Diepenbroek[4], Ethan Fast[6], Ragul Gowthaman[8,9], Jieling He[10], Vadim Karnaukhov[11,12], Dario F. Marzella[4], Pieter Meysman[13], Morten Nielsen[7], Jonas Birkelund Nilsson[7], Sebastian Nymann Deleuran[7], Farzaneh M. Parizi[14], Aurelien Pelissier[15,16], Brian G. Pierce[8,9], Maria Rodriguez Martinez[15], Dona Roran A R[4], Shayana Saravanakumar[8,9], Yanjun Shao[15], Nils Smit[4], Max Van Houcke[13], Gian Marco Visani[17], Yat-Tsai Richie Wan[7], Xiaowo Wang[10], Lawson Woods[3], Sander Wuyts[13], Chengkai Xiao[18], Li C. Xue[4], Justin Barton[19], Matthew Noakes[20], Damon H. May[20], Bjoern Peters[1,21*], IMMREP25 Participant Consortium

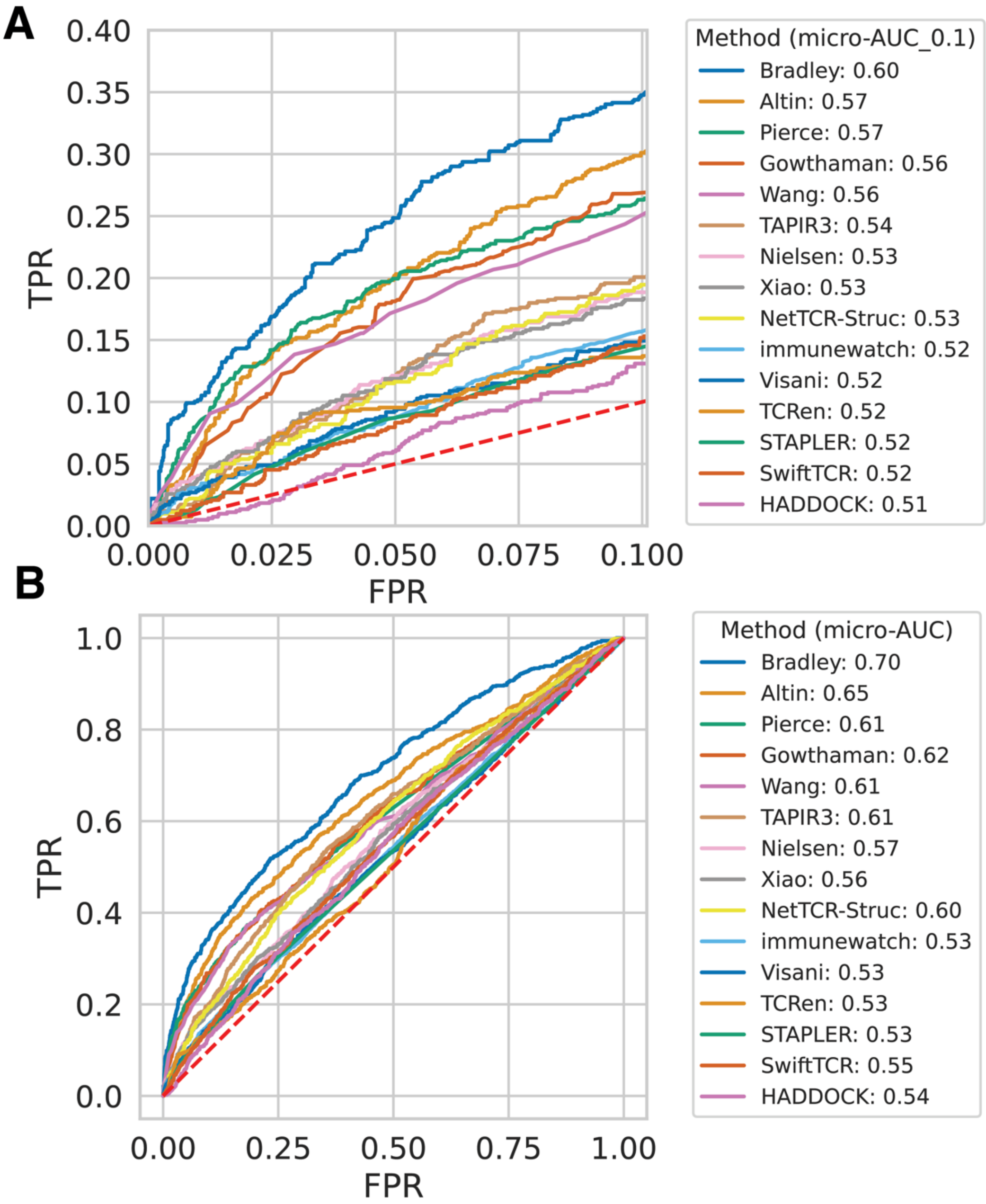


**Supplementary Figure 1:** micro-ROC curves and associated early retrieval AUCs (**B**) and full AUCs (**C, D**).

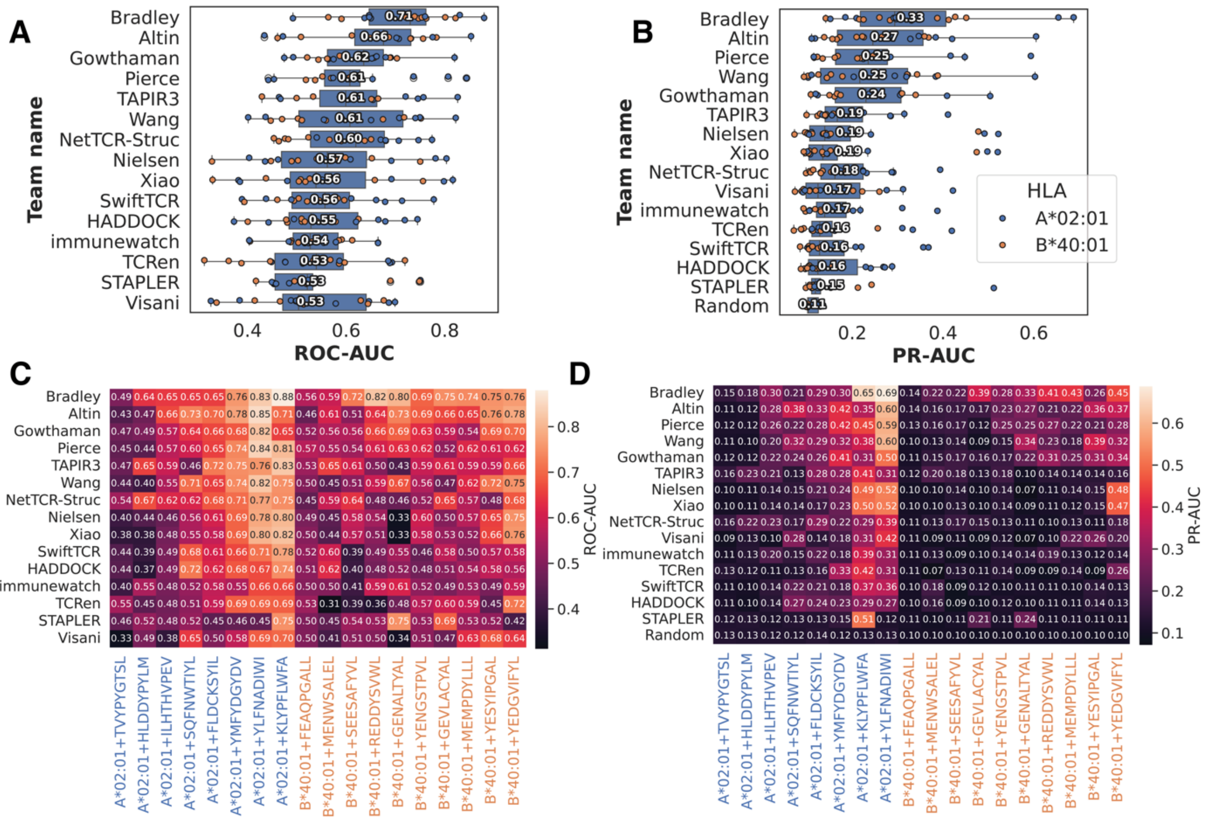


**Supplementary Figure 2:** ROC-AUC (**A**) and PR-AUC (**B**) ranking with mean values; these specific per-peptide values are shown in panels **C** and **D**. NB. that random performance varies by up to 4% per-peptide in the PR-AUC.

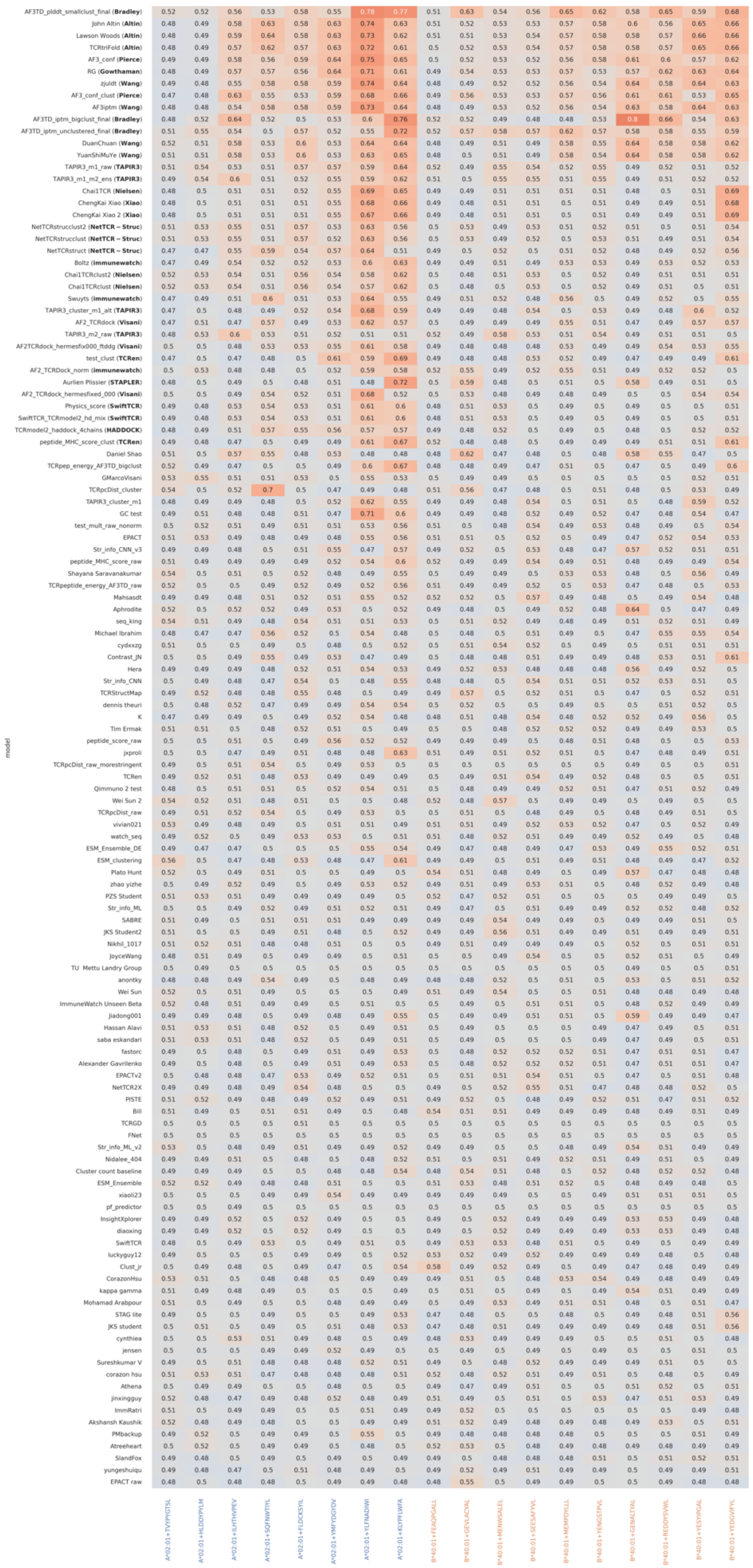
model
A*02:01+TVYPYGTSL
A*02:01+HLDDYPYLM
A*02:01+ILHTHVPEV
A*02:01+SQFNWTIYL
A*02:01+FLDCKSYIL
A*02:01+YMFYDGYDV
A*02:01+YLFNADIWI
A*02:01+KLYPFLWFA
B*40:01+FEAQPGALL
B*40:01+GEVLACYAL
B*40:01+MENWSALEL
B*40:01+SEESAFYVL
B*40:01+MEMPDYLLL
B*40:01+YENGSTPVL
B*40:01+GENALTYAL
B*40:01+REDDYSVWL
B*40:01+YESYIPGAL
B*40:01+YEDGVIFYL

**Supplementary Figure 3:** the full leaderboard with per-peptide ROC-AUC_0.1 values reported. The methods with macro-AUC_0.1 scores ≥ 0.52 are indicated by their associated team name in bold.

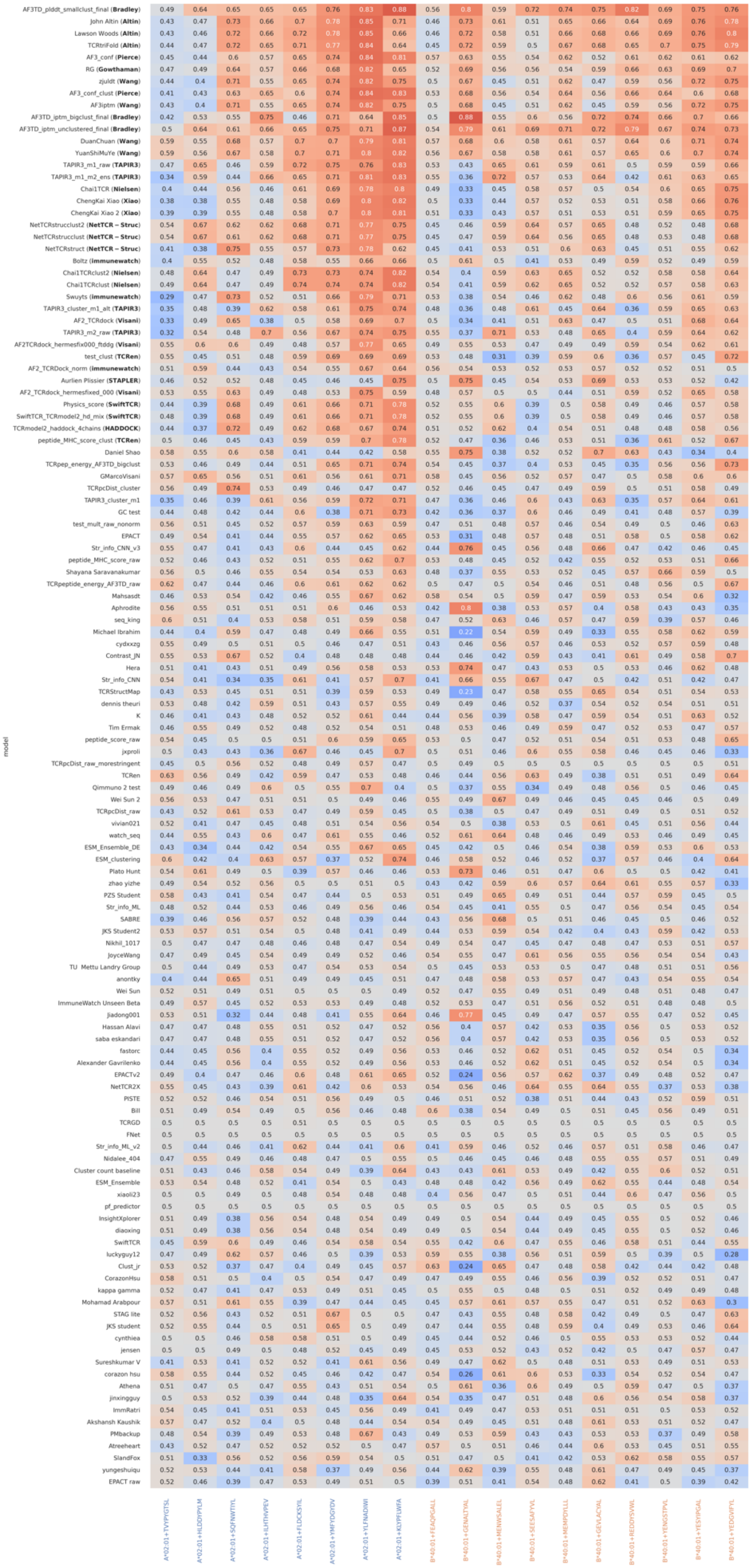
model

**Supplementary Figure 4:** the full leaderboard with per-peptide ROC-AUC values reported. The methods with macro-AUC_0.1 scores ≥ 0.52 are indicated by their associated team name in bold.

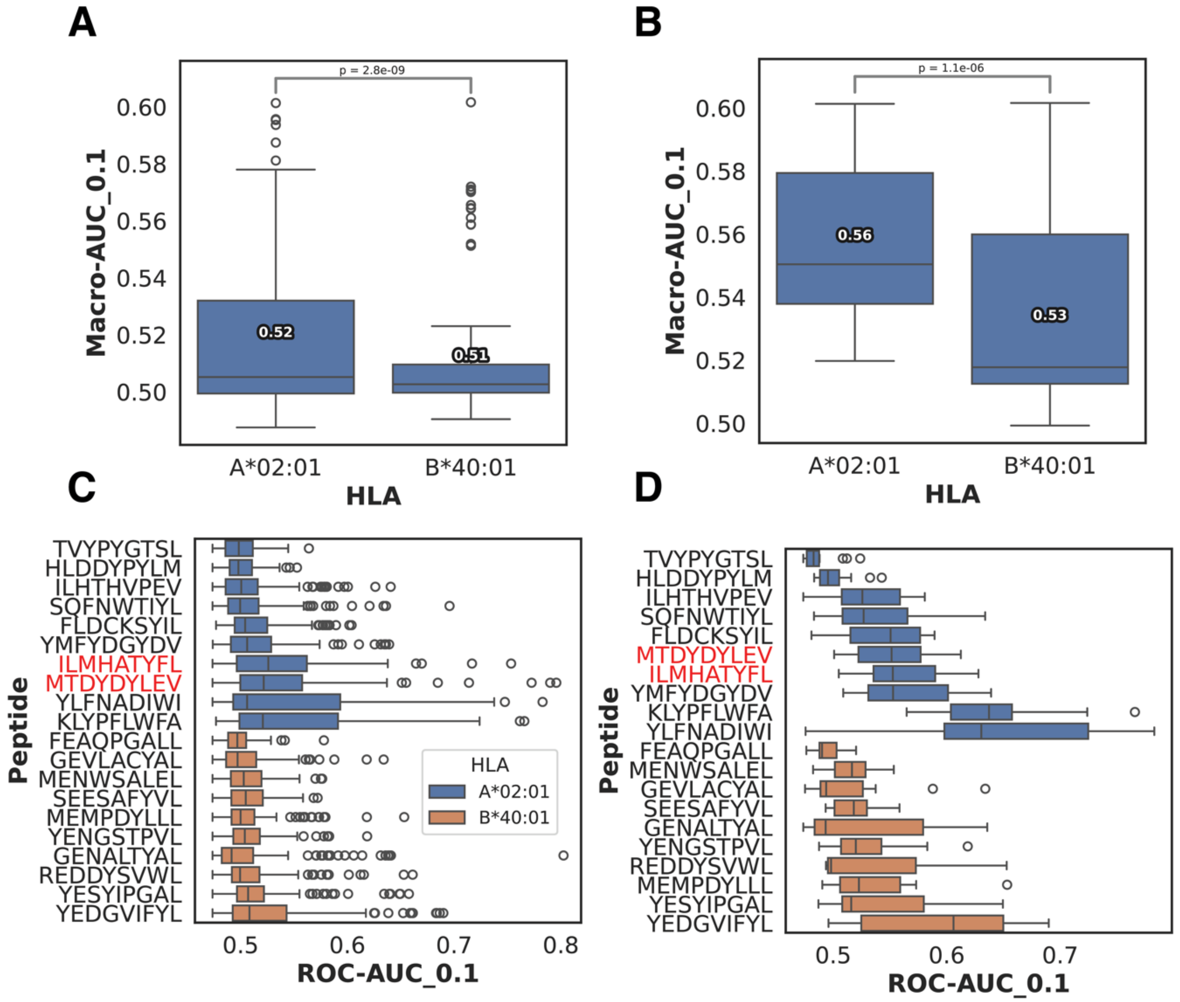


**Supplementary Figure 5:** variance across HLA and between peptides for all submissions (N = 126) (**A, C**) or for the teams which demonstrated significant predictive performance (**B, D**). Peptides in red are the public pMHCs (which are excluded from figures **A** and **B**).

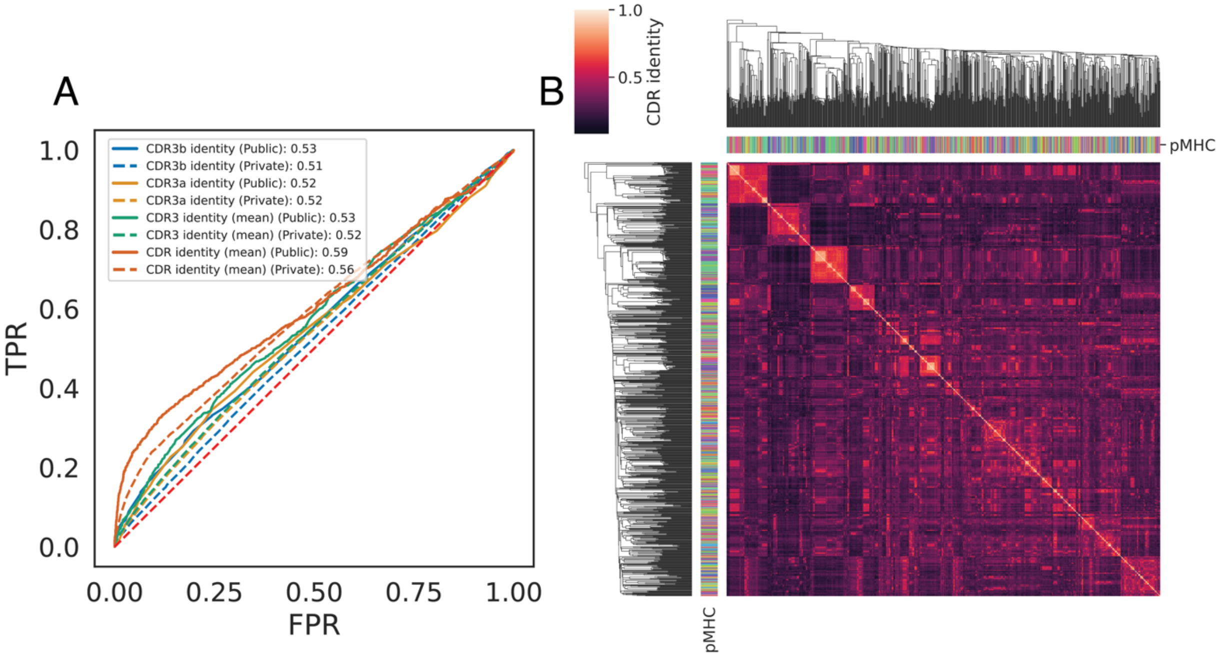


**Supplementary Figure 6:** the evaluation dataset has some underlying TCR sequence identity structure that weakly correlates with pMHC, e.g. TCRs sharing high CDR identity are likely to share the same pMHC (maximal ROC-AUC_0.1 of 0.59 for the public pMHC TCR set) (**A**). Panel **B** shows an average CDR identity clustering with rows colored by pMHC.

| Method | Macro AUC_0.1 |
|---|---|
| TAPIR3 | 0.538 |
| STAPLER | 0.518 |
| MixTCRpred1.0 | 0.510 |
| EPACT | 0.510 |
| NetTCR2.2 | 0.506 |
| MINT | 0.502 |
| TULIP | 0.500 |
| TEIM | 0.496 |

**Supplementary Table 1**: the top performing sequence-based model was TAPIR3; performance values are compared with published sequence-based methods.
